# How genetic disease risks can be misestimated across global populations

**DOI:** 10.1101/195768

**Authors:** Michelle S Kim, Kane P Patel, Andrew K Teng, Ali J Berens, Joseph Lachance

**Affiliations:** School of Biological Sciences, Georgia Institute of Technology

**Keywords:** ascertainment bias, genetic risk scores, genetic epidemiology, genome-wide association studies, global health, health disparities, population genetics

## Abstract

**Background:** Accurate assessment of health disparities requires unbiased knowledge of genetic risks in different populations. Unfortunately, most genome-wide association studies use genotyping arrays and European samples. Here, we integrate whole genome sequence data from global populations, results from thousands of GWAS, and extensive computer simulations to identify how genetic disease risks can be misestimated.

**Results:** In contrast to null expectations, we find that risk allele frequencies at known disease loci are significantly different for African populations compared to other continents. Strikingly, ancestral risk alleles are found at 9.51% higher frequency in Africa and derived risk alleles are found at 5.40% lower frequency in Africa. By simulating GWAS with different study populations, we find that non-African cohorts yield disease associations that have biased allele frequencies and that African cohorts yield disease associations that are relatively free of bias. We also find empirical evidence that genotyping arrays and SNP ascertainment bias contribute to continental differences in risk allele frequencies. Because of these causes, polygenic risk scores can be grossly misestimated for individuals of African descent. Importantly, continental differences in risk allele frequencies are only moderately reduced if GWAS use whole genome sequences and hundreds of thousands of cases and controls. Finally, comparisons between uncorrected and corrected genetic risk scores reveal the benefits of considering whether risk alleles are ancestral or derived.

**Conclusions:** Our results imply that caution must be taken when extrapolating GWAS results from one population to predict disease risks in another population.

## Background

In the past decade, over 3,500 genome-wide association studies (GWAS) have successfully identified more than 68,000 genetic associations with common diseases and other traits [1, 2]. However, the vast majority of published GWAS have used samples of European ancestry [3, 4], and a looming challenge is to be able to generalize GWAS results across populations [5-11]. An additional complication is that existing GWAS use genotyping arrays, as opposed to whole genome sequencing (WGS). Each disease-associated locus has risk and protective alleles, and results from GWAS can be combined to generate polygenic risk scores to predict individual risks of disease [12-14]. These polygenic risk scores quantify hereditary disease burdens by summing the number of risk alleles in each individual’s genome and sometimes weighting SNPs by effect size [15]. The “missing heritability” problem hampers genetic risk scores, as many causal variants remain undiscovered [16, 17]. Diseases can also have different genetic architectures in different populations [18]. Because of these issues, genetic predictions of disease risk are not always accurate, and it is important to be able to distinguish between situations where genetic risks actually differ between populations and when genetic predictions of differences in disease risks are spurious.

Although health disparities are often due to access to healthcare and socio-economic factors [19, 20], genetic differences in disease risks arise when allele frequencies at disease-associated loci differ across populations [15]. Populations that share recent ancestry have similar allele frequencies and hereditary disease risks, while populations that diverged in the deep past can have large allele frequency differences at disease-associated loci [21, 22]. These differences are magnified by population bottlenecks and founder effects, including elevated risks of cystic fibrosis among the Québécois [23] and cardiovascular disease among the descendants of the HMS Bounty mutineers [24]. However, many common diseases are polygenic [25, 26], and allele frequency differences at individual loci tend to average out. Because of this, the overall burden of hereditary disease is expected to be similar across the globe [27], with the possible exception of reduced genetic load in African populations [28]. For polygenic diseases, the null expectation is that individuals from different populations will have similar counts of risk alleles.

The genetic ancestry of study participants can cause hereditary disease risks to be misestimated. Indeed, genetic risk scores generated from different study cohorts have been shown to vary across populations [5]. As of 2016, the ancestry of 81% of all GWAS samples was European and 14% was Asian [3], and this is likely to cause the set of known disease associations to be enriched for alleles that are polymorphic or intermediate frequency in Europe or Asia, but not Africa. Inequities in genetic studies parallel what is observed in social science research: most samples are from <u>W</u>estern, <u>e</u>ducated, industrialized, rich and <u>d</u>emocratic (WEIRD) societies [29, 30]. For disease associations to be detected, loci need to be polymorphic in the study population. Because of this, disease loci with allele frequencies that are zero or one in European populations are likely to be missed (i.e. the “known unknowns” [31]), and some of these disease loci will have intermediate frequencies in other populations. Disease associations found in one population can over- or underestimate genetic disease risks in other populations. One partial solution to this problem is to perform multiethnic GWAS that include individuals from multiple populations [32].

Commonly used genotyping arrays can also cause predictions of hereditary disease risks to be misestimated. One issue is that SNPs on genotyping arrays tend to have large minor allele frequencies [33-35]. These older SNPs often have large allele frequency differences between populations [36, 37]. Systematic biases can also arise because commercially available genotyping arrays tend to use SNPs that were originally ascertained in European populations. This SNP ascertainment bias can be particularly problematic if it yields disease loci with risk allele frequencies that are high for one population and low for another population.

Demographic history also affects whether known disease associated loci have biased allele frequencies. Consider disease-associated alleles that are initially found at the same frequency in two populations, i.e. prior to divergence (Figure 1a). Note that risk alleles can be ancestral (shared with other primates) or derived (due to new mutations) and that ancestral alleles tend to be high frequency while derived alleles tend to be low frequency [38, 39]. Over time, allele frequencies at each locus diverge between daughter populations. Importantly, bottlenecked non-African populations have experienced greater amounts of genetic drift than African populations [40] (Figure 1b). This asymmetry, coupled with statistical power being maximized at intermediate allele frequencies [41], can cause known disease-associated loci to have biased allele frequencies. Specifically, we predict that non-African GWAS will catch disease loci that have higher ancestral risk allele frequencies (and lower derived allele frequencies) in Africa (Figure 1c). By contrast, we predict that African GWAS will catch a relatively unbiased set of disease-associated loci (Figure 1d). Although continental differences in ancestral and derived risk allele frequencies have been observed for prostate cancer loci [42], these biases have yet to be studied in a comprehensive way.

**Figure 1.**
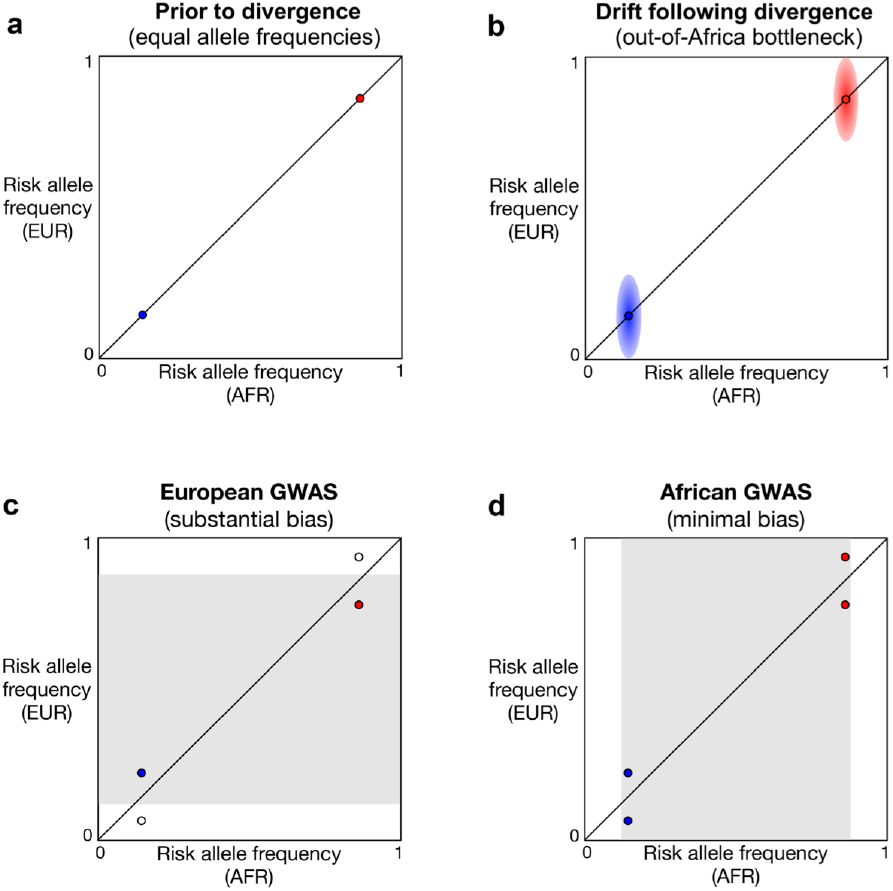
GWAS in bottlenecked European populations catch different types of disease loci than GWAS in non-bottlenecked African populations. Ancestral risk alleles are labelled red and derived risk alleles are labelled blue. Statistical power to detect associations is maximized at intermediate allele frequencies in the study population (gray shading). Filled circles indicate disease loci that are able to be caught by a GWAS, and open circles indicate disease loci that are unable to be caught by a GWAS. (**a**) Prior to divergence, allele frequencies are the same in both populations. (**b**) Non-African populations experience greater amounts of genetic drift. Diffusion of allele frequencies following divergence is indicated by red and blue shading. (**c**) European GWAS are predicted to catch derived risk alleles that have higher frequencies in Europe and ancestral risk alleles that have higher frequencies in Africa. (**d**) African GWAS are predicted to catch a relatively unbiased set of risk alleles.

At present, it is unknown how much the set of known disease associations hinders precision medicine and personal genomics. To bridge this knowledge gap, we integrated whole genome sequence data from global populations with results from thousands of GWAS and ran extensive computer simulations. These analyses: 1) revealed novel empirical patterns at disease-associated loci, 2) identified multiple causes of how disease risks can be misestimated in global populations, and 3) examined different solutions to this problem (including alternative GWAS study designs and building genetic risk scores that correct for major sources of bias).

## Results

### African risk allele frequencies differ from other continents

We tested whether there are any systematic biases in genetic estimates of disease risk by analyzing allele frequencies at 3036 GWAS loci for each continental population in the 1000 Genomes Project. Contrary to null expectations, mean risk allele frequencies are not the same for each population (Figure 2a). Overall, African populations have significantly higher risk allele frequencies compared to non-African populations (mean difference: +1.15%, p-value = 0.0213, paired Wilcoxon signed-rank test). Population-level differences in risk allele frequencies persist when disease associations are binned into seven different categories. Compared to other populations, African populations have the highest risk allele frequency for metabolic (p-value = 0.0055), morphological (p-value = 0.0949), cancer (p-value = 0.1169), neurological (p-value = 0.0995), and miscellaneous diseases (p-value = 0.3865, paired Wilcoxon signed-rank tests). African populations have intermediate frequencies of risk alleles at loci that are associated with GI or liver diseases (p-value = 0.6965), and lower frequencies of risk alleles at loci that are associated with cardiovascular disease (p-value = 0.0140, paired Wilcoxon signed-rank tests). These statistical comparisons reflect allele frequency differences at individual SNPs. Among non-African populations there is no underlying trend. Some of the continental patterns described here are at odds with clinical data (e.g. health disparities involving cardiovascular disease in African-Americans [43]). This discrepancy between clinical data and allele frequencies suggests that genetic disease risks may be misestimated for individuals with African ancestry.

**Figure 2.**
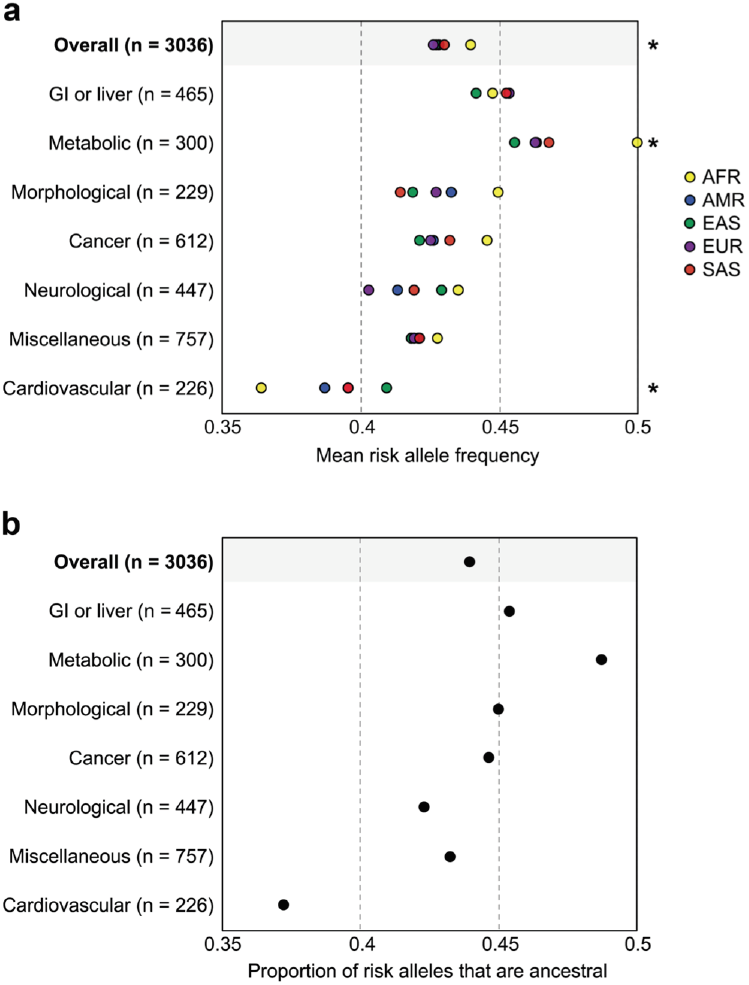
Known disease associations lead to misestimates of genetic disease risks. (**a**) Risk allele frequencies at published disease-associated loci from the NHGRI-EBI GWAS Catalog vary by population. * indicates a statistically significant allele frequency difference between African and non-African populations (p-values < 0.05, paired Wilcoxon rank sum tests). n = number of disease-associated loci per disease category. (**b**) Proportion of disease-associated loci where the risk allele is ancestral, as opposed to derived.

Disease categories that have a larger proportion of ancestral alleles tend to have elevated risk allele frequencies in Africa (Figure 2b). After binning GWAS loci by disease category, we find that the differences in the mean frequency of risk alleles between African and non-African populations are highly correlated with the proportion of risk alleles that are ancestral (r^2^ = 0.842). Accurate estimation of genetic disease risks across global populations may hinge upon knowledge of whether risk-increasing alleles are ancestral or derived.

### Ancestral and derived alleles yield different patterns of genetic disease risk

For loci that are not associated with any disease, the null expectation is that ancestral and derived allele frequencies will be broadly similar across global populations. Just because *Homo sapiens* emerged in Africa does not mean that African genomes have an excess of ancestral alleles – all human populations share the same evolutionary distance to chimpanzees. Due to the out-of-Africa bottleneck, African genomes are more likely to be heterozygous for derived alleles and non-African genomes are more likely to be homozygous for derived alleles. Examining WGS data from the 1000 Genomes Project, we find that derived allele frequencies (DAF) are similar for each population (Figure 3a). However, disease-associated loci need not exhibit the same pattern.

**Figure 3.**
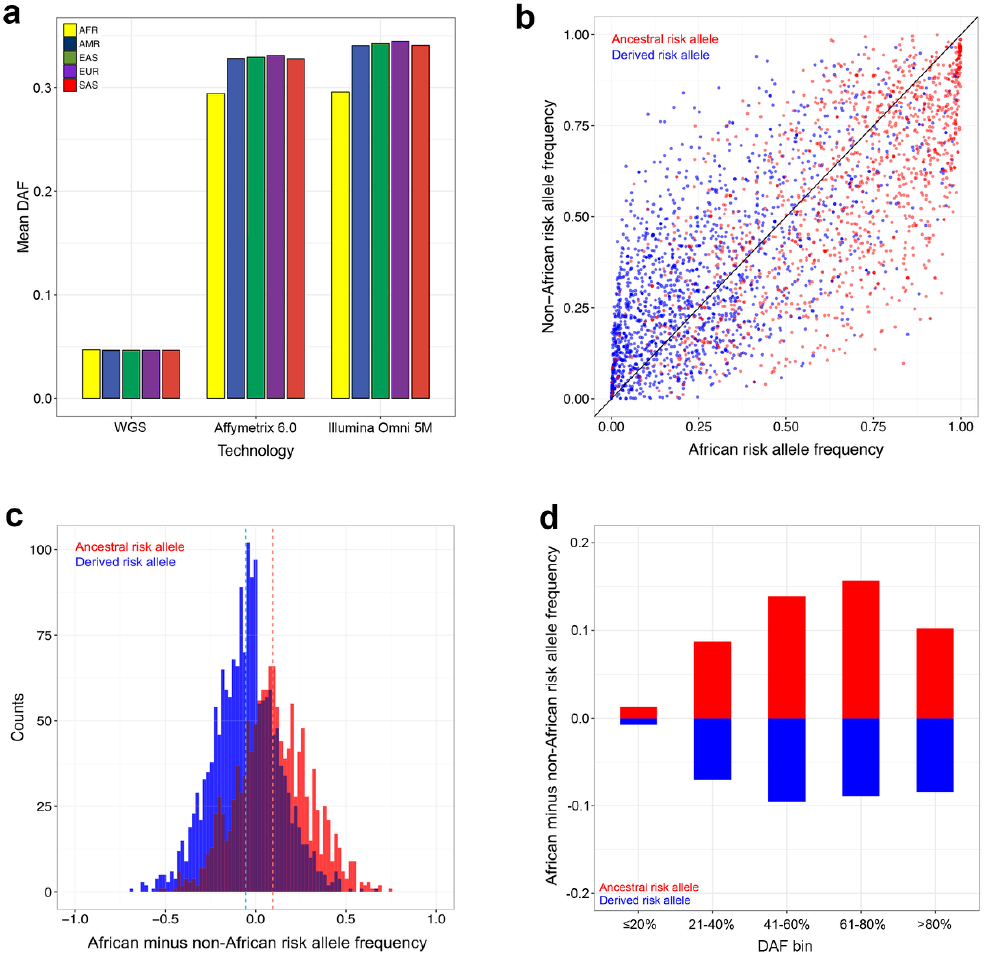
Empirical patterns depend on whether disease-associated alleles are ancestral or derived. (**a**) Mean derived allele frequencies of non-disease SNPs from whole genome sequencing and genotyping arrays. 1000 Genomes Project data are shown. (**b**) Joint SFS of published GWAS loci. Ancestral risk alleles are labelled red and derived risk alleles are labelled blue. (**c**) The frequencies of ancestral risk alleles are higher in Africa (+9.51% on average) and the frequencies of derived risk alleles are lower in Africa (−5.40% on average). Dashed lines indicate mean values. (**d**) Continental differences in risk allele frequencies are minimal for young SNPs. Disease-associated loci are binned by DAF and whether risk alleles are ancestral or derived.

The joint site frequency spectrum (SFS) enables the frequencies of individual risk alleles to be compared between African and non-African populations. Similar numbers of disease associations are found above and below the diagonal in Figure 3b. However, conditioning on whether risk alleles are ancestral or derived reveals a striking pattern: 69.2% of ancestral risk alleles are found at higher frequency in African populations (red dots below the diagonal), and 64.5% of derived risk alleles are found at higher frequency in non-African populations (blue dots above the diagonal). The magnitudes of allele frequency differences between populations also vary for ancestral and derived risk alleles. We find that ancestral risk alleles are found at much higher frequencies in Africa and derived risk alleles are found at moderately lower frequencies in Africa (Figure 3c). Specifically, the mean difference in ancestral risk allele frequencies between African and pooled non-African populations is +9.51%, and the mean difference in derived risk allele frequencies between African and pooled non-African populations is −5.40% (p-value < 2.2×10^-16^ for both comparisons, Wilcoxon signed-rank tests). The overall continental difference in risk allele frequencies of +1.15% arises because 44% of presently known disease-associated loci have ancestral risk alleles.

Derived allele frequencies serve as proxies for SNP age [44], and we find that older disease-associated loci are more likely to have large differences in continental allele frequencies. For each 20% DAF bin (pooled data), we calculated the difference in risk allele frequencies between African and non-African populations. In sharp contrast to other DAF bins, published disease loci with DAF ≤ 0.2 exhibit only a small amount of bias (Figure 3d). This pattern occurs regardless of whether risk alleles are ancestral or derived. Note that SNPs with DAF ≤ 0.2 tend to younger than 125,000 years old, assuming an effective population size of 10,000 individuals and generation times of 25 years [44].

### Choice of study population contributes to misestimates of genetic disease risk

Most disease associations have been discovered in study cohorts with European ancestry, and this can bias the estimation of genetic disease risks in diverse global populations. Empirical data reveal the effects of GWAS study populations: many disease-associated alleles segregate at intermediate frequencies in non-African populations but are found at extremely low or high frequencies in Africa (compare the vertical and horizontal borders of Figure 3b). This occurs because statistical power is maximized at intermediate frequencies, and most disease-associated loci have been discovered in non-African populations. Existing GWAS have discovered relatively few disease alleles that segregate only in African populations.

To further isolate the effects of different study populations we simulated a large number of GWAS results, varying the continental ancestry of each study cohort. Importantly, our GWAS simulations did not assume that there are any underlying differences in hereditary disease risks across populations. We find that computer simulations recapitulate empirical patterns at known disease loci, and that GWAS of bottlenecked non-African populations yield different results than GWAS of African populations (Figure 4). Simulated GWAS that use an African (AFR) cohort yield similar risk allele frequencies across each of the five continental populations. However, simulated GWAS that use American (AMR), East Asian (EAS), European (EUR), or South Asian (SAS) cohorts produce a set of disease-associated loci with elevated frequencies of ancestral risk alleles in Africa (Figure 4a) and reduced frequencies of derived risk alleles in Africa (Figure 4b). These simulation results indicate that systematic allele frequency differences between populations need not be due to any underlying difference in risk. The effects of European study cohorts are still seen when GWAS simulations use data from WGS, as opposed to genotyping arrays (Table 1). We also find that continental differences in risk allele frequencies occur if GWAS simulations use a more stringent p-value filter or simulations assume different modes of inheritance, including dominant or recessive disease alleles (Table S1 and Table S2). Additionally, GWAS simulations of study cohorts that contain a mixture of individuals from different populations still yield disease-associated loci with continental biases in risk allele frequencies (MIX in Figure 4). These results suggest that pooling samples with different ancestries is unlikely to completely alleviate the problem of misestimating genetic disease risks. Regardless of the choice of study cohort, allele frequencies are similar for each non-African population, reflecting the relatively recent divergence times between these populations.

**Table 1.** Differences in allele frequencies between African and European populations for different genotyping technologies

| Data type | Allele frequency difference between Africa and Europe |  |
| --- | --- | --- |
|  | Ancestral risk allele | Derived risk allele |
| NHGRI-EBI GWAS Catalog (empirical) | +11.7% | -6.7% |
| Affymetrix Genome-Wide Human SNP Array 6.0 (simulated*) | +10.7% | -8.0% |
| Illumina Omni 5M microarray (simulated*) | +11.0% | -8.2% |
| Whole genome sequences (simulated*) | +9.7% | -7.2% |

\*GWAS simulation parameters: sample size = 3500 cases and 3500 controls, study population = EUR, p-value threshold = $1 \times 10^{-5}$ , mode of inheritance = additive, prevalence = 0.1, genotype relative risk = 1.211.

**Figure 4.**
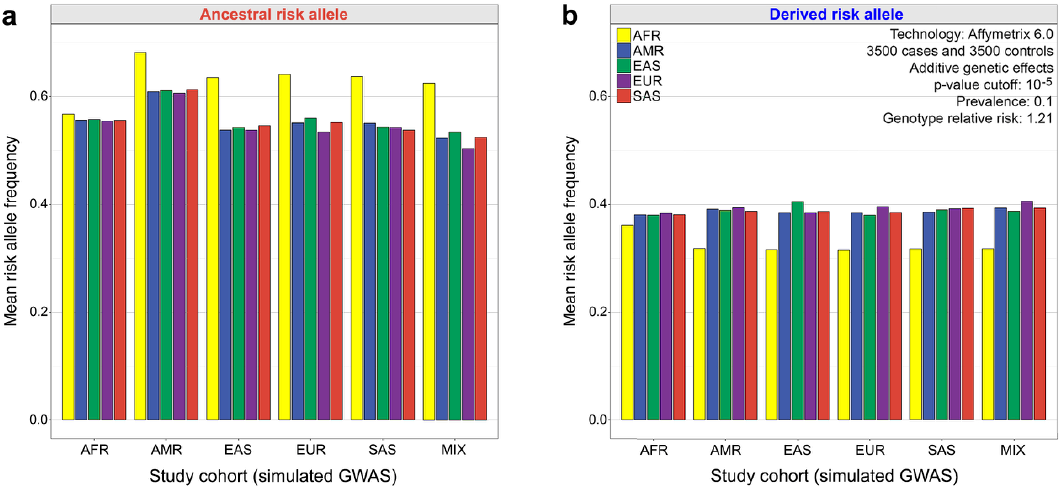
GWAS simulations reveal the effects of different study cohorts. Mean risk allele frequencies in different continental populations are shown for each study cohort (3036 disease associations per simulation). Despite the absence of any underlying differences in risk, disease-associated loci that that are detected in non-African study cohorts have biased frequencies. (**a**) GWAS simulations where the ancestral allele increases risk. (**b**) GWAS simulations where the derived allele increases risk.

We also examined the effects of genotype-by-environment (GxE) interactions by allowing effect sizes to vary by population in our GWAS simulations. In general, results from these simulations mirror results other simulations: ancestral risk allele frequencies are higher in African populations than non-African populations, and derived risk allele frequencies are lower in African populations than non-African populations (Figure S2). Compared to African study cohorts, European study cohorts magnify these allele frequency differences between populations. Choice of study cohort imposes a filter on effect sizes, as SNPs with very small effect sizes do not yield detectable associations (compare gray pre-GWAS effects sizes to red and blue post-GWAS effect sizes in Figures S2-S4). Large effect sizes enable high frequency ancestral alleles and low frequency derived alleles to be detected in a GWAS. The results described above are also robust to systematic biases in effect sizes, i.e. scenarios where pre-GWAS European effect sizes tend to be larger than African effect-sizes or vice versa (Figures S3 and S4).

### Genotyping arrays and SNP ascertainment bias cause disease risks to be misestimated

Many commonly used genotyping arrays contain SNPs that were ascertained in a relatively small number of European individuals. This ascertainment bias results in allele frequency distributions that vary by genotyping platform. Compared to WGS data, derived allele frequencies are higher for SNPs on the Affymetrix Genome-Wide Human SNP Array 6.0 and the Illumina Omni 5M microarray. SNPs on genotyping arrays also exhibit continental biases (Figure 3a). Specifically, we find that derived allele frequencies in African populations are markedly lower than derived allele frequencies in non-African populations (p-value < 2.2×10^-16^ for both arrays, Wilcoxon signed-rank tests).

The joint SFS of non-African and African populations further reveals the effects of SNP ascertainment bias. Examining WGS data, we find that similar numbers of SNPs have elevated derived allele frequencies in non-African and African populations (Figure S1a). By contrast, the Affymetrix Genome-Wide Human SNP Array 6.0 and the Illumina Omni 5M microarray are enriched SNPs with higher derived allele frequencies outside of Africa (i.e. SNPs above the diagonal in Figure S1b and Figure S1c). Importantly, this pattern mirrors what is seen for empirical GWAS data (Figure S1d), which suggests that genotyping arrays contribute to continental differences in risk allele frequencies at known disease-associated loci.

Because many disease-associations involve imputed SNPs, we also tested whether continental differences in risk allele frequencies persist for disease-associated loci that are not on the Affymetrix Genome-Wide Human SNP 6.0 Array. For this empirical set of disease-associated loci, we find that sites with ancestral risk alleles have higher allele frequencies in Africa (+8.63% on average) and that SNPs with derived risk alleles have lower allele frequencies in Africa (−4.83% on average). This suggests that biases persist even for imputed SNPs.

### Continental differences in allele frequencies persist even if whole genome sequencing and large sample sizes are used

Simulations of GWAS results were used to infer the extent that misestimates of disease risks depend upon genotyping technology (Table 1). Here, simulations assume European ancestry for each study cohort and sample sizes of 3500 cases and 3500 controls. We find that different genotyping arrays yield similar results: the Affymetrix Genome-Wide Human SNP Array 6.0 and the Illumina Omni 5M microarray yield ancestral risk allele frequencies that are 10.7% and 11.0% higher in Africa and derived risk alleles that are 8.0% and 8.2% higher in Europe, respectively. Somewhat surprisingly, continental differences in allele frequencies also occur for GWAS simulations that use WGS data. Focusing on WGS GWAS simulations, ancestral risk allele frequencies are 9.7% higher in Africa and derived risk alleles are 7.2% higher in Europe. These patterns arise because of our choice of study cohort and because sample sizes of 3500 cases and 3500 controls have relatively little power to catch rare disease alleles.

Continental biases in risk allele frequencies occur even if GWAS use large sample sizes. Simulated GWAS with less than 10,000 European cases and controls yield large differences in African and non-African allele frequencies (Figure 5). This occurs regardless of whether simulations use SNPs from the genotyping arrays or WGS. Increasing GWAS sample sizes increases the statistical power to detect associations with rare alleles. However, our simulations reveal that there are diminishing returns for increasing sample sizes, especially if GWAS use genotyping arrays. Well-powered studies with hundreds and thousands of cases and controls still yield notable differences in continental allele frequencies – even if WGS are used (Figure 5). These results indicate that WGS is unable to completely mitigate the effects different study populations.

**Figure 5.**
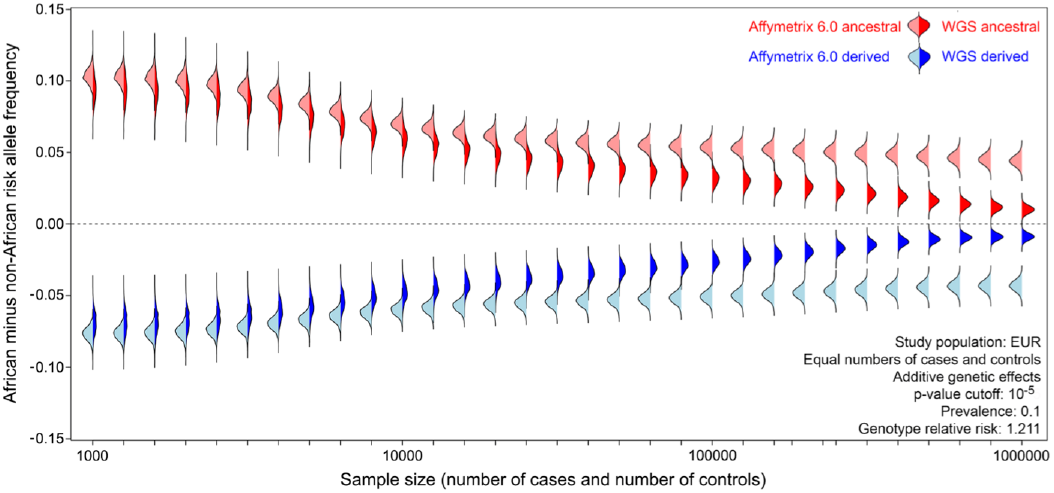
GWAS simulations reveal that continental differences in allele frequencies persist even if whole genome sequencing and large sample sizes are used. Bean plots show the results of 1000 simulations per set of parameter values (3036 disease associations per simulation). Simulations using SNPs on genotyping arrays are represented by light shading and simulations using WGS data are represented by dark shading. Colors indicate whether risk alleles are ancestral (red) or derived (blue). Sample sizes shown are the number of cases and the number of controls.

### Correcting for ancestral and derived risk alleles leads to improved genetic risk scores

Standardized genetic risk scores (GRS) were generated for 2504 individuals and seven different disease categories. This involved integrating a curated list of disease-associated loci from the NHGRI-EBI GWAS Catalog with individual-level genotype data from the 1000 Genomes Project. Positive GRS values indicate genomes that contain more risk alleles than the global mean, and negative GRS values indicate genomes that contain less risk alleles than the global mean. Standardized GRS are scaled in terms of standard deviations from the mean, i.e. they are Z-scores. In general, different populations have GRS distributions that mirror what is seen for allele frequency data (compare Figure 6 to Figure 2a). We find that African individuals have uncorrected GRS that differ from other populations (p-value = 0.0037 for GI or liver diseases and p-value < 2.2×10^-16^ for all other disease categories, Mann-Whitney U tests). These differences are larger for metabolic, cancer, and cardiovascular disease risks. There is a substantial amount of overlap between the GRS distributions of each non-African population, and this pattern occurs for all disease categories. Within each population there is also a large range of GRS values. Also note that admixed genomes from the Americas (AMR in Figure 6) have GRS that are broadly similar to other non-African genomes. Although GRS reflect an individual’s genetic propensity for different disease categories, we caution against over-interpreting these results. This is because GRS have been built from a biased set of disease-associated loci.

**Figure 6.**
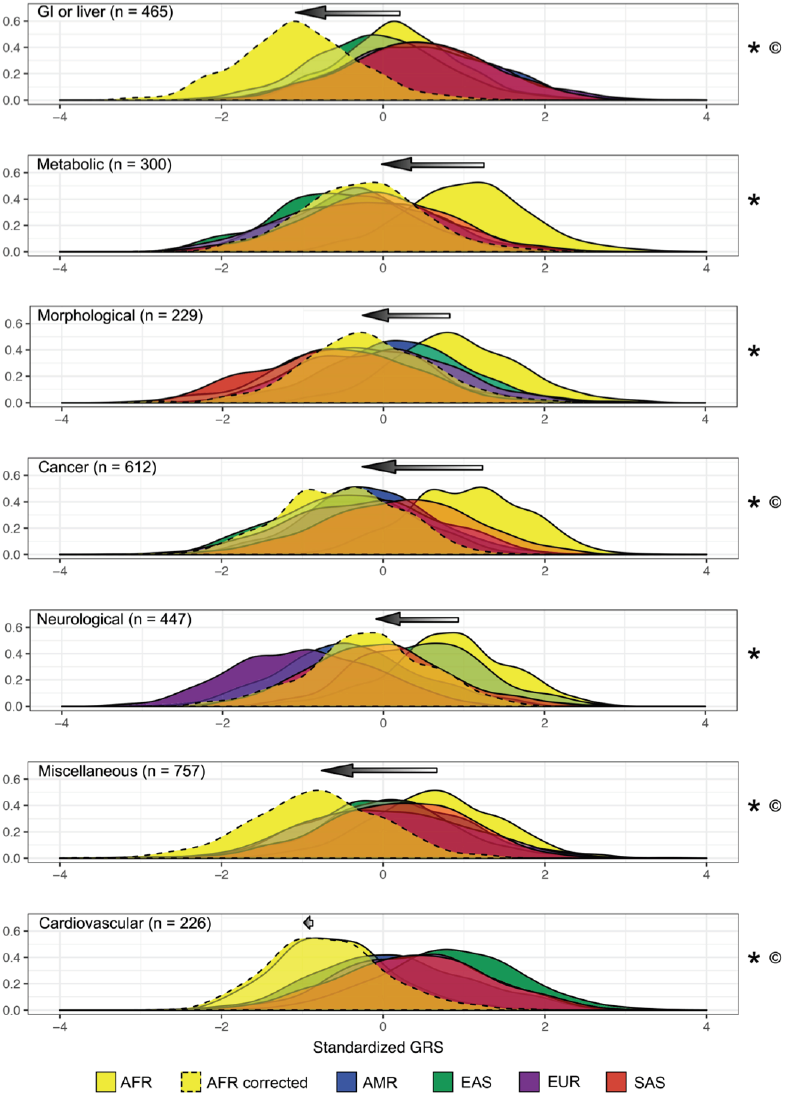
Genetic risk scores (GRS) before and after correcting for ancestral and derived risk alleles. GRS probability densities for each continental population are shown (solid lines: uncorrected GRS, dashed lines: corrected GRS for African genomes). n = number of disease-associated loci per disease category. Arrows indicate the shift in African GRS after correcting for whether risk alleles are ancestral or derived. * indicates uncorrected African GRS that are significantly different than non-African GRS, and © indicates corrected African GRS that are significantly different than non-African GRS (p-values < 0.05, Mann Whitney U tests).

GRS corrections reduce some, but not all, of the population-level differences in predicted disease risks. Here, we compensate for continental differences in ancestral and derived risk allele frequencies by generating corrected GRS for African genomes. We find that African individuals have corrected GRS that are similar to other populations for metabolic (p-value = 0.8080), morphological (p-value = 0.0671), and neurological disease risks (p-value = 0.7116, Mann-Whitney U tests). By contrast, African individuals have corrected GRS that are different than other populations for GI or liver, cancer, miscellaneous, and cardiovascular disease risks (p-value < 2.2×10^-16^ for each disease category, Mann-Whitney U tests). Corrections involve in a leftward shift in the GRS of African genomes, the magnitude of which depends on the proportion of ancestral risk alleles for each disease category (compare the size of arrows in Figure 6). We observe three different outcomes: minimal effects, over-correction, and reduction of bias. Cardiovascular risk predictions for African genomes were largely unchanged (i.e. GRS still appear to underestimate the risks of cardiovascular disease in individuals of African descent). Two disease categories (GI or liver and miscellaneous diseases) have corrected GRS distributions that differ more between African and non-African populations than uncorrected GRS distributions. The remaining four disease categories (metabolic, morphological, cancer, and neurological diseases) have corrected GRS distributions that overlap heavily with other populations. Although the correction method used here alleviates some forms of bias, our results suggest that GRS can be further improved by considering additional parameters.

## Discussion

The biased set of disease associations that are presently known causes hereditary disease risks to be misestimated. Specifically, African populations tend to have higher frequencies of ancestral risk alleles and lower frequencies of derived risk alleles at existing GWAS loci. Considering the magnitude of these differences and the proportion of disease-associated alleles that are ancestral, as opposed to derived, yields risk allele frequencies that are 1.15% higher in Africa. Elevated risk allele frequencies in African populations are the opposite of what one expects to see given human demographic history. Due to population bottlenecks, non-African populations are expected to have greater amounts of genetic load [28]. This discrepancy arises because GWAS rely on European study cohorts and data from genotyping arrays. Continental differences in allele frequencies also have important ramifications for precision medicine and personal genomics: disease risks are likely to be misestimated if GWAS results are naively used to calculate genetic risk scores (Figure 6). This can obscure existing health disparities that are due to socio-cultural factors including access to medical care [45, 46]. High risk individuals may have genetic profiles that lull them into a false sense of security, and low risk individuals may have genetic risk profiles that lead to an undue amount of worry.

Here, we are concerned with the limitations of using disease associations discovered in one population to predict disease risks in another population, as opposed to whether GWAS findings can be successfully replicated across multiple populations. The effects of different study cohorts are asymmetric. Non-African GWAS results can be used to predict disease risks in other non-African populations, but these disease associations generalize poorly to African populations (Figure 4). By contrast, African GWAS results can be used to predict disease risks in a relatively unbiased way across all global populations. This asymmetry arises as a by-product of demographic history and the out-of-Africa migration (Figure 1) and because GWAS use arrays that suffer from SNP ascertainment bias (Figure 3a). Our results suggest that there may be additional benefits to including a large number of African individuals in multiethnic GWAS. We note that difficulties can still arise when transferring GWAS results from one non-African population to another non-African population. This is due to both the existence of private risk alleles and divergence times that can exceed 30,000 years. Regardless of the study cohort used to generate genetic risk scores, it is impossible to fully correct for missing risk alleles from understudied populations. Problems generalizing GWAS results cannot be solved by only using WGS and large sample sizes (Figure 5). Furthermore, many variants discovered by WGS are rare and population-specific. That said, genetic risk scores generated from WGS data are expected to be less biased than genetic risk scores generated from array data, especially when sample sizes are large.

Although this paper focuses on risk allele frequency differences across populations, we note that many disease loci remain undetected, and this also contributes to misestimates of disease risks. These missing disease loci are particularly important when risk alleles are population-specific. This underscores the need for genetic epidemiology studies to include samples from a diverse set of populations.

Our study demonstrates the benefits of adopting an evolutionary perspective towards health and disease [47, 48]. Important empirical patterns would not have been noticed without considering ancestral vs. derived states of alleles. Continental differences in allele frequencies also depend upon SNP age. An evolutionary perspective is also valuable for understanding how genetic disease risks can be misestimated across populations. Specifically, we find that it matters whether populations have experienced a history of bottlenecks and founder effects. Knowing whether individual disease loci have experienced a history of natural section can lead to additional insights [42, 49, 50]. That said, systematic allele frequency biases can be mistaken for directional selection, hindering tests of polygenic selection acting on GWAS traits [51].

Recently, Martin et al. found that polygenic risk scores yield inaccurate predictions of height and schizophrenia, and that GRS for Type II Diabetes depend upon on choice of study cohort [5]. Using coalescent simulations, they also found that the proportion of heritability that can be explained decreases with distance to the GWAS study population. Using complementary approaches, our study resulted in novel discoveries. We find that ancestral and derived states of risk alleles play a central role in the estimation of genetic disease risks across multiple populations, something missed by prior studies that examine the generalizability of GWAS results. We also find that important asymmetries exist when extrapolating results between African and non-African populations, and that population bottlenecks play a key role (i.e. generalizability of results depends on more than the evolutionary distance between populations). By explicitly testing the effects of different genotyping technologies and sample sizes, we were able to discover that WGS of hundreds of thousands of cases and controls still yields biased GWAS results. Martin et al. also advocate mean-centering GRS for each population [5], but this solution can be problematic if hereditary disease risks actually differ between populations.

Our GRS calculations illustrate how misestimation of genetic risks can obscure whether there are any real differences in disease risks across populations (Figure 6). Two types of error are possible: 1) The underlying risk of a particular disease may actually be the same for different populations, yet GRS distributions show little overlap. 2) The underlying risk of a particular disease may actually differ for populations, yet GRS distributions show extensive overlap. Accurate GRS corrections are needed to exclude either of these two possibilities. Environmental effects and genotype-by-environment interactions also contribute to disease phenotypes [52]. Studies of immigrants, admixed families, and adopted individuals may prove to be particularly informative with respect to genetics and health inequities [53-56]. Corrected GRS for admixed genomes may also benefit from the use of local ancestry painting tools like RFMix [57] or ELAI [58].

## Conclusions

Going forward, multiple approaches can be used to extend the benefits of precision medicine and personal genomics to a wide range of global populations. One option is to assume that disease associations can be generalized across populations without any complications. However, this approach is flawed because only a biased set of disease loci are known at present. A second option is to require that genetic risk scores only use disease associations discovered in the same population (i.e. avoid generalizing results across populations). However, this is unfeasible from a logistical standpoint – as it would require repeating every GWAS in every global population. A third option is to use whole genome sequencing and large African study cohorts to generate sets of disease associated-loci that can be generalized relatively free of bias. On a more practical side, genetic risk scores can be generated that correct for existing biases. This requires understanding how risk allele frequencies differ between populations (as shown here) and leveraging linkage disequilibrium information to infer the effect sizes of risk alleles in non-study populations [59, 60]. Finally, we note that the gold-standard for evaluating the genetic risk scores involves testing how well they predict disease phenotypes in diverse populations – something that requires individual-level phenotype data. Only by understanding population genetics and the effects of SNP ascertainment bias can accurate predictive models of genetic disease risks be built.

## Methods

### Population genetic data

Allele frequencies were obtained for each of the five continental populations of the 1000 Genomes Project: Africa (AFR), Americas (AMR), East Asia (EAS), Europe (EUR), and South Asia (SAS) [21]. These frequencies were used to generate risk allele frequencies and derived allele frequencies at disease-associated loci from the NHGRI-EBI GWAS Catalog and simulated datasets. Ancestral and derived states in phase 3 1000 Genomes Project VCF files were used (these ancestral states were inferred via the EPO pipeline from Ensembl). We found that derived allele frequencies in all populations were elevated for large chunks of chromosome 8, which is indicative of misidentified ancestral states. To compensate for this, we masked SNPs found in the chr8: 89,000,000-146,364,022 region (hg19). Individuals in phase 3 of 1000 Genomes Project were genotyped using WGS. Allele frequencies of SNPs on the Affymetrix Genome-Wide Human SNP Array 6.0 and the Illumina Omni 5M microarray were found by merging data from the 1000 Genomes Project with lists of SNP IDs obtained from the Affymetrix and Illumina websites.

### Identification of disease-associated variants

Using the NHGRI-EBI GWAS Catalog [1], Berens et al. generated a curated set of 3180 disease-associated loci [61]. This involved filtering out SNPs that were not associated with a disease, eliminating SNPs lacking risk allele or odds ratio information, and LD-pruning. Here, we further constrained the set of disease-associated loci from [61] by requiring knowledge of whether risk alleles are ancestral or derived. After excluding 144 SNPs with unknown ancestral states, we were left with a focal set of 3036 disease-associated loci (Table S3). We classified these 3036 disease-associated loci into seven non-overlapping categories: gastrointestinal/liver, metabolic, morphological, cancer, neurological, miscellaneous, and cardiovascular. Wilcoxon signed-rank tests were used to compare disease allele frequencies between African and non-African populations. Disease-associated loci were binned by DAF, averaging across all 1000 Genomes Populations. Allele ages were estimated as per Eq. 4 in [44] (assuming N=10,000 and a generation time of 25 years).

### GWAS simulations

Computer simulations were used to test whether SNP ascertainment bias alone can produce what appears to be genetic differences in disease risks across populations. The goal here was to generate simulated datasets comparable to the set of 3036 disease-associated loci from the NHGRI-EBI GWAS Catalog. These simulations assume that the underlying risks of disease are the same across the globe. Two general types of simulations were run: simulations with ancestral risk alleles and simulations with derived risk alleles. Simulations involved randomly drawing a *test SNP* from a list of known genetic variants ascertained via WGS or found on commercial genotyping arrays. Conditioning on whether risk alleles are ancestral or derived, the risk allele frequency of the *test SNP* was found in the study population. We then used a Perl script based on the GAS/CaTS power calculator [41] to determine the probability of detecting a successful genetic association at the *test SNP*. The GAS power calculator leverages information about the number of cases and controls, p-value threshold, disease model, prevalence, disease allele frequency, and genotype relevant risk (http://csg.sph.umich.edu/abecasis/cats/gas_power_calculator/). For each *test SNP*, we generated a uniformly distributed random number between 0 and 1. The *test SNP* was retained if the random number was less than the power to successfully detect a genetic association, and the *test SNP* was rejected if the random number was greater than the probability of detection. This process was repeated until a set of 3036 successful disease associations were detected. At each of these 3036 SNPs, we obtained simulated risk allele frequencies for five populations in the 1000 Genomes Project dataset (AFR, AMR, EAS, EUR, SAS). Our default parameters were as follows: genotyping technology = Affymetrix Genome-Wide Human SNP Array 6.0, study population = Europe (EUR), sample size = 3500 cases and 3500 controls, genetic model = additive, p-value threshold = 10^-5^, prevalence = 0.1, and genotype relative risk = 1.211. These parameter values were chosen to be representative of the empirical data found in the NHGRI-EBI GWAS Catalog.

Our default model was modified to test which aspects of SNP ascertainment bias contribute the most to continental differences in risk allele frequencies. This involved varying the following simulation parameters: genotyping technology, sample size, mode of inheritance, and the p-value threshold required for association detection. To examine the effects of different study populations, simulated risk allele frequencies were chosen from one of five different populations (AFR, AMR, EAS, EUR, or SAS) or from an equal mixture of all five populations (MIX). The effects of different sample sizes were simulated by varying the number of cases and controls from 3 to 6 on a log10 scale at intervals of 0.1 (i.e. between 1,000 and 1,000,000 cases and controls). The effects of different genotyping technologies were simulated by drawing random SNPs from either the Affymetrix Genome-Wide Human SNP Array 6.0, the Illumina Omni 5M microarray, or WGS data from the 1000 Genomes Project. Three genetic modes of inheritance were simulated: dominant, additive, and recessive. Two different p-value thresholds were simulated: 1×10^-5^ and 5×10^-8^.

We also simulated the results of GWAS where effect sizes vary between populations. Simulations examined three different effect size distributions (symmetric, larger effect sizes in Europe, and larger effect sizes in Africa), two different types of risk alleles (ancestral and derived), and two different study cohorts (European and African). In each simulation run, 3036 disease-associated loci were obtained using the power calculator described above. Simulations were repeated 1000 times per combination of parameters. Symmetric effect sizes were generated by drawing locus-specific genotype relative risks for each *test SNP* from a gamma distribution (shape= 1.24, scale = 0.85). These parameter values were chosen to give a distribution of effect sizes that is comparable to loci in the NHGRI-EBI GWAS Catalog. We allowed genotype relative risks for each *test SNP* to vary by population by adding random noise (normally distributed, mean = 0, standard deviation = 0.5). Simulated genotype relative risks <1 were set equal to 1. Larger European effect sizes were generated by drawing locus-specific genotype relative risks from a gamma distribution that was shifted 0.5 upwards (Figure S3). Larger African effect sizes were generated by drawing locus-specific genotype relative risks from a gamma distribution shifted 0.5 to the right (Figure S4). A representative dataset from GWAS simulations is included in Table S4

### GRS corrections

Genetic risk scores (GRS) for 2504 individuals were built using genotypes at a curated set of 3036 disease-associated loci from the NHGRI-EBI GWAS Catalog. Note that genetic risk scores are sometimes called polygenic risk scores (PRS). For each disease locus we counted whether an individual has 0, 1, or 2 copies of the risk allele. Because each disease category includes a heterogeneous set of diseases and phenotypes we did not incorporate odds ratio and/or effect size information into our GRS calculations. Counts of risk alleles were then summed across all loci that belong to a particular disease category, yielding a raw GRS for each individual. Standardized GRS values were calculated for each combination of individual and disease category by finding the mean and standard deviation of raw GRS values across all 2504 individuals in the 1000 Genomes Project dataset. Given our empirical results (Figure 3c), diploid African genomes tend to have 0.1902 (2 x 9.51%) additional copies each ancestral risk allele and 0.1082 (2 x 5.41%) fewer copies of each derived risk allele compared to non-African genomes. Because of this, our correction method considered the state of the risk alleles (ancestral or derived). Uncorrected African GRS use counts of 0,1, or 2 risk alleles at each disease locus. Corrected African GRS use counts of −0.1902, 0.8098, and 1.8098 “effective risk alleles” for ancestral alleles and 0.1082, 1.1082, and 2.1082 “effective risk alleles” for derived alleles. The same mapping of raw GRS to standardized GRS was used for uncorrected and corrected African GRS.

## Additional files

**Table S1.** Effects of different p-value thresholds for GWAS simulations.

**Table S2.** GWAS simulations of dominant, additive, and recessive disease alleles.

**Table S3.** Curated set of 3036 disease-associated loci from the NHGRI-EBI GWAS Catalog.

**Table S4.** Representative set of loci from GWAS simulations

**Figure S1.** Joint site frequency spectra for multiple genotyping technologies.

**Figure S2.** GWAS simulations that allow effect sizes to vary by population.

**Figure S3.** GWAS simulations with larger effect sizes in Europe.

**Figure S4.** GWAS simulations with larger effect sizes in Africa.

## Acknowledgements

We thank A. Martin, U. Martinez-Marigorta, M. Quiver, and C. Simonti for helpful discussions during the writing of this paper.

## Funding

This work was supported by NIH/NCI grant U01CA184374 and start-up funds from Georgia Institute of Technology.

## Availability of data and materials

Global allele frequencies are publicly available from the 1000 Genomes Project website: http://www.internationalgenome.org/data. Disease associations are publicly available from the NHGRI-EBI GWAS Catalog: https://www.ebi.ac.uk/gwas. R and Perl scripts used in GWAS simulations are available from GitHub at https://github.com/LachanceLab/AscertainmentBias_GWAS.

## Authors’ contributions

MSK performed statistical analyses and GWAS simulations, generated genetic risk scores, and wrote the manuscript. KPP analyzed population genetic data. AKT generated an initial set of genetic risk scores. AJB curated the set of disease-associated loci. JL conceived and supervised the study, interpreted results, and wrote the manuscript. All authors have read and approved the final manuscript.

## Competing interests

All authors declare that they have no competing interests.

## Supplementary tables

**Table S1.** Effects of different p-value thresholds for GWAS simulations.

| P-value threshold | Allele frequency difference<br>between Africa and Europe |  |
| --- | --- | --- |
|  | Ancestral<br>risk allele | Derived<br>risk allele |
| $1 \times 10^{-5}$ | +10.7% | -8.0% |
| $5 \times 10^{-8}$ | +12.2% | -8.8% |

GWAS simulation parameters: technology = Affymetrix Genome-Wide Human SNP Array 6.0, sample size = 3500 cases and 3500 controls, study population = EUR, mode of inheritance = additive, prevalence = 0.1, genotype relative risk = 1.211.

**Table S2.** GWAS simulations of dominant, additive, and recessive disease alleles.

| Mode of inheritance | Allele frequency difference<br>between Africa and Europe |  |
| --- | --- | --- |
|  | Ancestral<br>risk allele | Derived<br>risk allele |
| Dominant | +19.7% | -2.2% |
| Additive | +10.7% | -8.0% |
| Recessive | +2.9% | -17.9% |

GWAS simulation parameters: technology = Affymetrix Genome-Wide Human SNP Array 6.0, sample size = 3500 cases and 3500 controls, study population = EUR, p-value threshold = $1 \times 10^{-5}$ , prevalence = 0.1, genotype relative risk = 1.211.

Table S3. Curated set of 3036 disease-associated loci from the NHGRI-EBI GWAS Catalog.

[S3_curated_disease_loci.txt]

Table S4. Representative set of loci from GWAS simulations. Simulation parameters: technology = Affymetrix Genome-Wide Human SNP Array 6.0, sample size = 3500 cases and 3500 controls, mode of inheritance = additive genetic effects, p-value cutoff = 10^-5^, prevalence = 10%, study cohort = EUR, genotype relative risks = vary symmetrically across populatios,

[Table S4_representative_loci_from_simulations.txt]

## Supplementary figures

**Figure S1.**
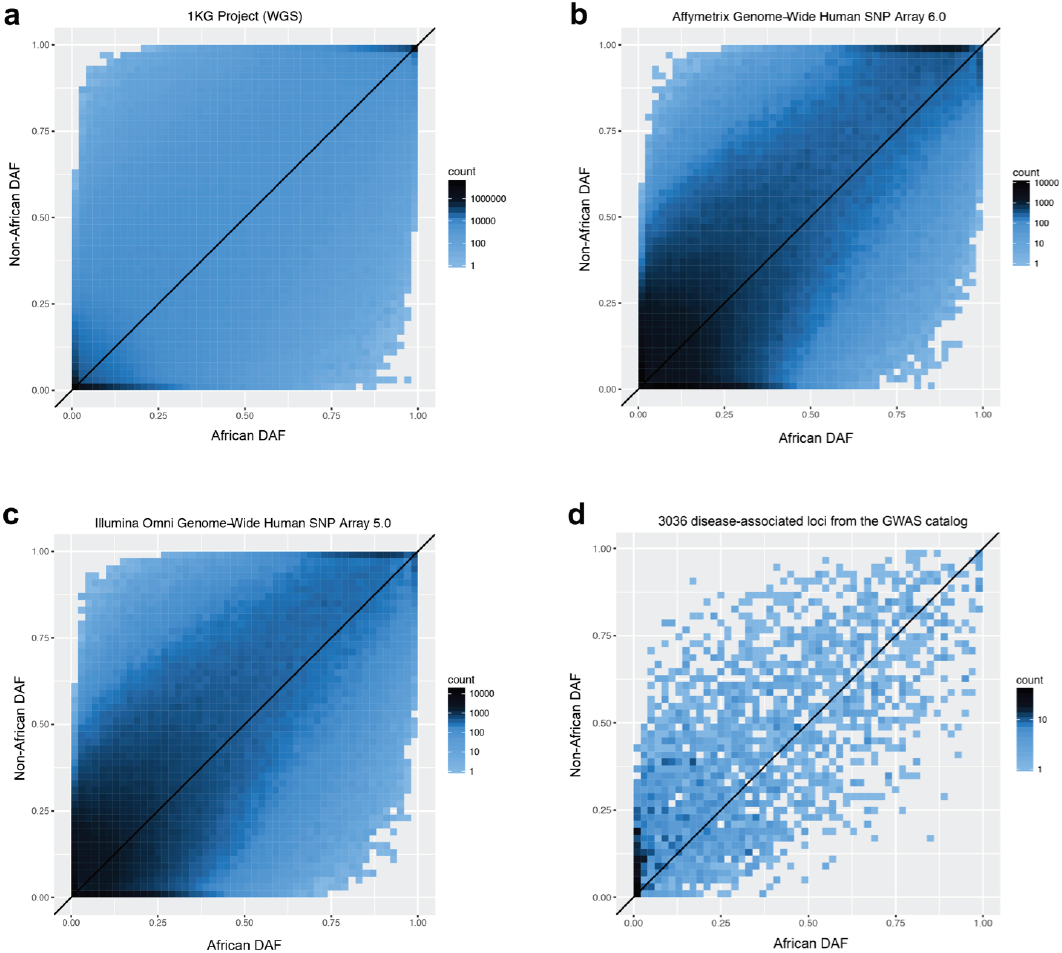
Empirical Joint site frequency spectra of for multiple genotyping technologies. DAF are shown in each panel. (**a**) Joint SFS of whole genome sequence (WGS) data. Non-African and African data from the 1000 Genomes Project are shown. Shading indicates counts of SNPs. (**b**) Joint SFS of ascertained SNPs on the Affymetrix Genome-Wide Human SNP Array 6.0. (**c**) Joint SFS of ascertained SNPs on the Illumina Omni 5M microarray. (**d**) Joint SFS of published disease-associated loci from the NHGRI-EBI GWAS Catalog.

**Figure S2.**
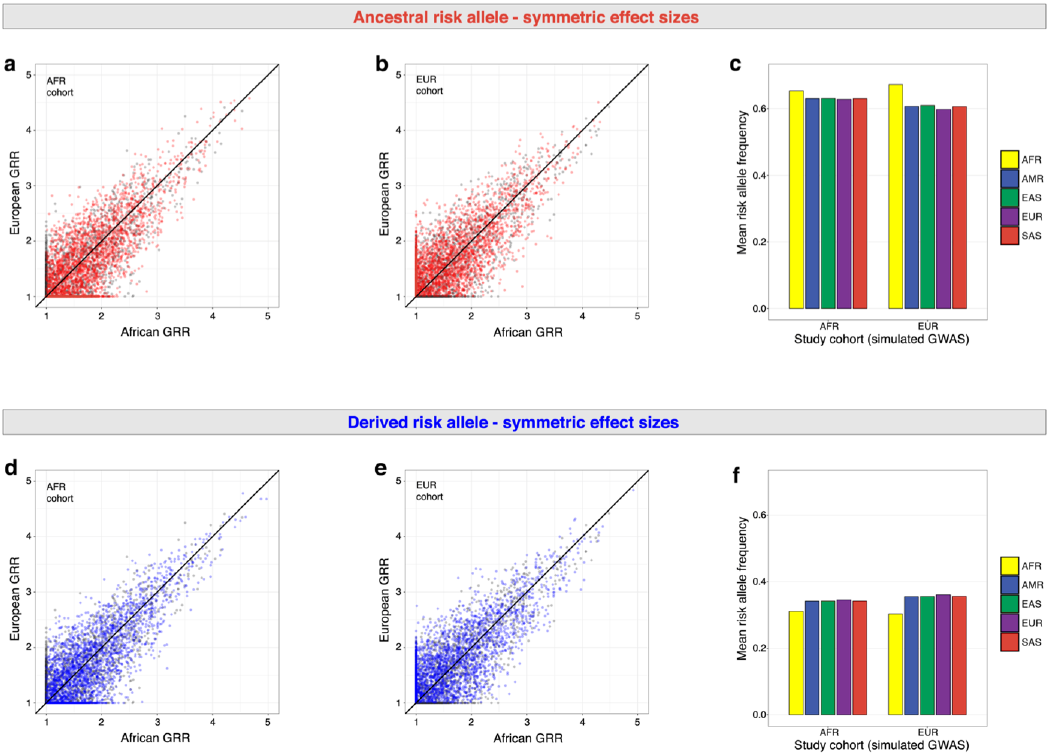
GWAS simulations that allow effect sizes to vary by population. Simulation parameters: technology = Affymetrix Genome-Wide Human SNP Array 6.0, sample size = 3500 cases and 3500 controls, mode of inheritance = additive genetic effects, p-value cutoff = 10^-5^, prevalence = 10%. Representative sets of 3036 SNPs are shown. Panels (**a**), (**b**), and (**c**) show the results of GWAS simulations where the ancestral allele increases risk. Panels (**d**), (**e**), and (**f**) show the results of GWAS simulations where the derived allele increases risk. Panels (**a**), (**b**), (**d**) and (**e**) show representative effect sizes in Europe and Africa, where GRR refers to genotype relative risk. Pre-GWAS effect sizes are indicated by gray points. Post-GWAS effect sizes are indicated by red points (ancestral risk alleles) and by blue points (derived risk alleles). Prior to GWAS simulations, effect sizes are symmetric. Mean risk allele frequencies in different continental populations are shown for each study cohort in panels (**c**) and (**f**).

**Figure S3.**
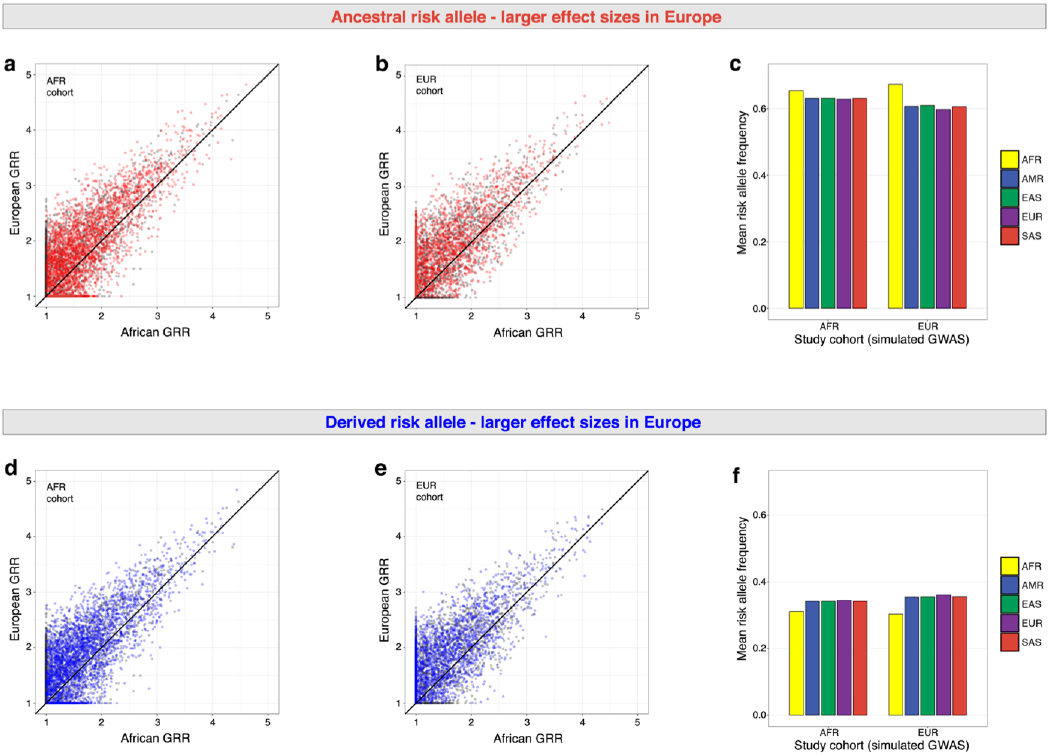
GWAS simulations with larger effect sizes in Europe. Simulation parameters are the same as in Figure S2, with the exception of larger GRR values in Europe. Representative sets of 3036 SNPs are shown. Panels (**a**), (**b**), and (**c**) show the results of GWAS simulations where the ancestral allele increases risk. Panels (**d**), (**e**), and (**f**) show the results of GWAS simulations where the derived allele increases risk. Panels (**a**), (**b**), (**d**) and (**e**) show representative effect sizes in Europe and Africa, where GRR refers to genotype relative risk. Pre-GWAS effect sizes are indicated by gray points. Post-GWAS effect sizes are indicated by red points (ancestral risk alleles) and by blue points (derived risk alleles). Prior to GWAS simulations, effect sizes are shifted upwards (i.e. higher in Europe). Mean risk allele frequencies in different continental populations are shown for each study cohort in panels (**c**) and (**f**).

**Figure S4.**
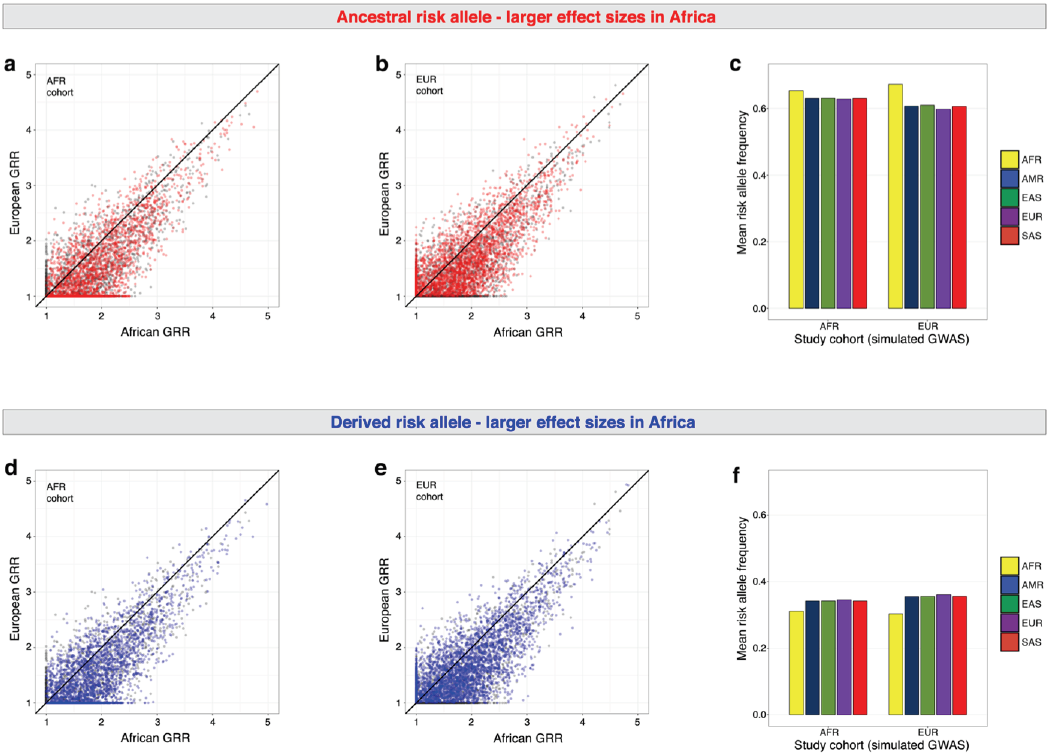
GWAS simulations with larger effect sizes in Africa. Simulation parameters are the same as in Figure S2, with the exception of larger GRR values in Africa. Panels (**a**), (**b**), and (**c**) show the results of GWAS simulations where the ancestral allele increases risk. Representative sets of 3036 SNPs are shown. Panels (**d**), (**e**), and (**f**) show the results of GWAS simulations where the derived allele increases risk. Panels (**a**), (**b**), (**d**) and (**e**) show representative effect sizes in Europe and Africa, where GRR refers to genotype relative risk. Pre-GWAS effect sizes are indicated by gray points. Post-GWAS effect sizes are indicated by red points (ancestral risk alleles) and by blue points (derived risk alleles). Prior to GWAS simulations, effect sizes are shifted to the right (i.e. larger in Africa). Mean risk allele frequencies in different continental populations are shown for each study cohort in panels (**c**) and (**f**).

## References

1. MacArthur J, Bowler E, Cerezo M, Gil L, Hall P, Hastings E, Junkins H, McMahon A, Milano A, Morales J, et al: The new NHGRI-EBI Catalog of published genome-wide association studies (GWAS Catalog). Nucleic Acids Res 2017, 45:D896–D901.

2. Hindorff LA, Sethupathy P, Junkins HA, Ramos EM, Mehta JP, Collins FS, Manolio TA: Potential etiologic and functional implications of genome-wide association loci for human diseases and traits. Proc Natl Acad Sci U S A 2009, 106:9362–9367.

3. Popejoy AB, Fullerton SM: Genomics is failing on diversity. Nature 2016, 538:161– 164.

4. Manolio TA: In Retrospect: A decade of shared genomic associations. Nature 2017, 546:360–361.

5. Martin AR, Gignoux CR, Walters RK, Wojcik GL, Neale BM, Gravel S, Daly MJ, Bustamante CD, Kenny EE: Human Demographic History Impacts Genetic Risk Prediction across Diverse Populations. The American Journal of Human Genetics 2017, 100:635–649.

6. Bustamante CD, Burchard EG, De la Vega FM: Genomics for the world. Nature 2011, 475:163–165.

7. Marigorta UM, Navarro A: High trans-ethnic replicability of GWAS results implies common causal variants. PLoS Genet 2013, 9:e1003566.

8. Palmer C, Pe’er I: Statistical correction of the Winner’s Curse explains replication variability in quantitative trait genome-wide association studies. PLoS Genet 2017, 13:e1006916.

9. Shriner D: Mixed Ancestry and Disease Risk Transferability. Current Genetic Medicine Reports 2015, 3:151–157.

10. Coram MA, Fang H, Candille SI, Assimes TL, Tang H: Leveraging Multi-Ethnic Evidence for Risk Assessment of Quantitative Traits in Minority Populations. Am J Hum Genet 2017.

11. Hindorff LA, Bonham VL, Brody LC, Ginoza MEC, Hutter CM, Manolio TA, Green ED: Prioritizing diversity in human genomics research. Nat Rev Genet 2018, 19:175–185.

12. Chatterjee N, Shi J, Garcia-Closas M: Developing and evaluating polygenic risk prediction models for stratified disease prevention. Nat Rev Genet 2016, 17:392–406.

13. International Schizophrenia Consortium, Purcell SM, Wray NR, Stone JL, Visscher PM, O’Donovan MC, Sullivan PF, Sklar P: Common polygenic variation contributes to risk of schizophrenia and bipolar disorder. Nature 2009, 460:748–752.

14. Shi J, Park JH, Duan J, Berndt ST, Moy W, Yu K, Song L, Wheeler W, Hua X, Silverman D, et al: Winner’s Curse Correction and Variable Thresholding Improve Performance of Polygenic Risk Modeling Based on Genome-Wide Association Study Summary-Level Data. PLoS Genet 2016, 12:e1006493.

15. Corona E, Chen R, Sikora M, Morgan AA, Patel CJ, Ramesh A, Bustamante CD, Butte AJ: Analysis of the genetic basis of disease in the context of worldwide human relationships and migration. PLoS Genet 2013, 9:e1003447.

16. Manolio TA, Collins FS, Cox NJ, Goldstein DB, Hindorff LA, Hunter DJ, McCarthy MI, Ramos EM, Cardon LR, Chakravarti A, et al: Finding the missing heritability of complex diseases. Nature 2009, 461:747–753.

17. Wray NR, Yang J, Hayes BJ, Price AL, Goddard ME, Visscher PM: Pitfalls of predicting complex traits from SNPs. Nat Rev Genet 2013, 14:507–515.

18. McClellan J, King MC: Genetic heterogeneity in human disease. Cell 2010, 141:210–217.

19. Warnecke RB, Oh A, Breen N, Gehlert S, Paskett E, Tucker KL, Lurie N, Rebbeck T, Goodwin J, Flack J: Approaching health disparities from a population perspective: the National Institutes of Health Centers for Population Health and Health Disparities. American journal of public health 2008, 98:1608–1615.

20. Woolf SH, Braveman P: Where health disparities begin: the role of social and economic determinants--and why current policies may make matters worse. Health Aff (Millwood) 2011, 30:1852–1859.

21. 1000 Genomes Project Consortium: A global reference for human genetic variation. Nature 2015, 526:68–74.

22. Li JZ, Absher DM, Tang H, Southwick AM, Casto AM, Ramachandran S, Cann HM, Barsh GS, Feldman M, Cavalli-Sforza LL, Myers RM: Worldwide human relationships inferred from genome-wide patterns of variation. Science 2008, 319:1100–1104.

23. Laberge AM, Michaud J, Richter A, Lemyre E, Lambert M, Brais B, Mitchell GA: Population history and its impact on medical genetics in Quebec. Clin Genet 2005, 68:287–301.

24. Macgregor S, Bellis C, Lea RA, Cox H, Dyer T, Blangero J, Visscher PM, Griffiths LR: Legacy of mutiny on the Bounty: founder effect and admixture on Norfolk Island. Eur J Hum Genet 2010, 18:67–72.

25. Timpson NJ, Greenwood CMT, Soranzo N, Lawson DJ, Richards JB: Genetic architecture: the shape of the genetic contribution to human traits and disease. Nat Rev Genet 2018, 19:110–124.

26. Visscher PM, Wray NR, Zhang Q, Sklar P, McCarthy MI, Brown MA, Yang J: 10 Years of GWAS Discovery: Biology, Function, and Translation. Am J Hum Genet 2017, 101:5–22.

27. Lohmueller KE: The distribution of deleterious genetic variation in human populations. Curr Opin Genet Dev 2014, 29:139–146.

28. Henn BM, Botigue LR, Peischl S, Dupanloup I, Lipatov M, Maples BK, Martin AR, Musharoff S, Cann H, Snyder MP, et al: Distance from sub-Saharan Africa predicts mutational load in diverse human genomes. Proc Natl Acad Sci U S A 2016, 113:E440–449.

29. Jones D: A WEIRD View of Human Nature Skews Psychologists’ Studies. Science 2010, 328:1627–1627.

30. Henrich J, Heine SJ, Norenzayan A: Most people are not WEIRD. Nature 2010, 466:29–29.

31. Logan DC: Known knowns, known unknowns, unknown unknowns and the propagation of scientific enquiry. J Exp Bot 2009, 60:712–714.

32. Pulit SL, Voight BF, de Bakker PI: Multiethnic genetic association studies improve power for locus discovery. PLoS One 2010, 5:e12600.

33. Clark AG, Hubisz MJ, Bustamante CD, Williamson SH, Nielsen R: Ascertainment bias in studies of human genome-wide polymorphism. Genome Res 2005, 15:1496– 1502.

34. McCarthy MI, Abecasis GR, Cardon LR, Goldstein DB, Little J, Ioannidis JP, Hirschhorn JN: Genome-wide association studies for complex traits: consensus, uncertainty and challenges. Nat Rev Genet 2008, 9:356–369.

35. Nielsen R: Population genetic analysis of ascertained SNP data. Hum Genomics 2004, 1:218–224.

36. Lachance J, Tishkoff SA: SNP ascertainment bias in population genetic analyses: why it is important, and how to correct it. Bioessays 2013, 35:780–786.

37. Albrechtsen A, Nielsen FC, Nielsen R: Ascertainment biases in SNP chips affect measures of population divergence. Mol Biol Evol 2010, 27:2534–2547.

38. Lachance J: Disease-associated alleles in genome-wide association studies are enriched for derived low frequency alleles relative to HapMap and neutral expectations. BMC Med Genomics 2010, 3:57.

39. Di Rienzo A, Hudson RR: An evolutionary framework for common diseases: the ancestral-susceptibility model. Trends Genet 2005, 21:596–601.

40. Ramachandran S, Deshpande O, Roseman CC, Rosenberg NA, Feldman MW, Cavalli-Sforza LL: Support from the relationship of genetic and geographic distance in human populations for a serial founder effect originating in Africa. Proc Natl Acad Sci U S A 2005, 102:15942–15947.

41. Skol AD, Scott LJ, Abecasis GR, Boehnke M: Joint analysis is more efficient than replication-based analysis for two-stage genome-wide association studies. Nat Genet 2006, 38:209–213.

42. Lachance J, Berens AJ, Hansen MEB, Teng AK, Tishkoff SA, Rebbeck TR: Genetic hitchhiking and population bottlenecks contribute to prostate cancer disparities in men of African descent. Cancer Res 2018, 78:2432–2443.

43. Benjamin EJ, Virani SS, Callaway CW, Chamberlain AM, Chang AR, Cheng S, Chiuve SE, Cushman M, Delling FN, Deo R: Heart disease and stroke statistics—2018 update: a report from the American Heart Association. Circulation 2018, 137:e67– e492.

44. Slatkin M, Rannala B: Estimating allele age. Annu Rev Genomics Hum Genet 2000, 1:225–249.

45. Braveman P, Egerter S, Williams DR: The social determinants of health: coming of age. Annu Rev Public Health 2011, 32:381–398.

46. Manrai AK, Funke BH, Rehm HL, Olesen MS, Maron BA, Szolovits P, Margulies DM, Loscalzo J, Kohane IS: Genetic Misdiagnoses and the Potential for Health Disparities. N Engl J Med 2016, 375:655–665.

47. Stearns SC, Medzhitov R: Evolutionary medicine. Sunderland, Massachussetts: Sinauer Associates, Inc., Publishers; 2016.

48. Crespi BJ: The emergence of human-evolutionary medical genomics. Evol Appl 2011, 4:292–314.

49. Bigham AW, Magnaye K, Dunn DM, Weiss RB, Bamshad M: Complex signatures of natural selection at GYPA. Human genetics 2018, 137:151–160.

50. Shriner D, Rotimi CN: Whole genome sequence-based haplotypes reveal single origin of the sickle allele during the Holocene Wet Phase. Am J Hum Genet 2018, 102:547–556.

51. Novembre J, Barton NH: Tread Lightly Interpreting Polygenic Tests of Selection. Genetics 2018, 208:1351–1355.

52. Hunter DJ: Gene-environment interactions in human diseases. Nat Rev Genet 2005, 6:287–298.

53. Hemminki K, Bermejo JL, Försti A: Opinion: The balance between heritable and environmental aetiology of human disease. Nature Reviews Genetics 2006, 7:958.

54. Haugaard JJ, Hazan C: Adoption as a natural experiment. Dev Psychopathol 2003, 15:909–926.

55. Sankar P, Cho MK, Condit CM, Hunt LM, Koenig B, Marshall P, Lee SS, Spicer P: Genetic research and health disparities. JAMA 2004, 291:2985–2989.

56. Fine MJ, Ibrahim SA, Thomas SB: The role of race and genetics in health disparities research. Am J Public Health 2005, 95:2125–2128.

57. Maples BK, Gravel S, Kenny EE, Bustamante CD: RFMix: a discriminative modeling approach for rapid and robust local-ancestry inference. Am J Hum Genet 2013, 93:278–288.

58. Guan Y: Detecting structure of haplotypes and local ancestry. Genetics 2014, 196:625–642.

59. Vilhjalmsson BJ, Yang J, Finucane HK, Gusev A, Lindstrom S, Ripke S, Genovese G, Loh PR, Bhatia G, Do R, et al: Modeling Linkage Disequilibrium Increases Accuracy of Polygenic Risk Scores. Am J Hum Genet 2015, 97:576–592.

60. Rosenberg NA, Huang L, Jewett EM, Szpiech ZA, Jankovic I, Boehnke M: Genome-wide association studies in diverse populations. Nat Rev Genet 2010, 11:356–366.

61. Berens AJ, Cooper TL, Lachance J: The genomic health of ancient hominins. Human Biology 2017, 89:5–17.

